# Pathogenic impact of ABCA4 missense variants in the structurally uncharacterized ECD1 region: implications for Stargardt disease

**DOI:** 10.64898/2026.06.25.734683

**Authors:** Nadee N. J. Matarage Don, Subhasis B. Biswas, Esther E. Biswas-Fiss

## Abstract

Pathogenic mutations in the *ABCA4* gene cause several inherited retinal diseases, particularly Stargardt disease (STGD1). However, many missense variants remain classified as variants of uncertain significance (VUS) due to inconclusive evidence regarding their pathogenic impact. The missense VUS span across all the domains of ABCA4, with the majority found in the larger extracellular domains (ECDs). The largest uncharacterized region of ABCA4 is located in ECD1, where limited structural information and inconsistent computational predictions hinder clinical interpretation of missense VUS in this region. Here, we integrated *in silico* analysis with in vitro functional assays to evaluate the pathogenicity of VUS in this region and improve their diagnostic classification. Missense VUS in the ECD1 uncharacterized region were curated from ClinVar. Six multiallelic sites were identified in the uncharacterized region and 13 missense VUS on these multiallelic sites were characterized using the integrated analysis. In the *in silico* platform, the pathogenicity of the VUS were predicted using multiple algorithms, and the structural effects of the variants were analyzed compared to the wild type. Recombinant variants were expressed in virus-like particles (VLPs), and protein expression, membrane localization, and ATPase activity were quantified relative to wild type to identify potential disease-causing variants.

From the integrated analysis, variants with pronounced structural destabilization, impaired membrane trafficking, and reduced or absent N-retinylidene-phosphatidylethanolamine (NRPE) substrate stimulated ATPase activities were identified as potentially deleterious. Notably, VUS at p.H193P and p.I214N showed loss of function, with p.I214N reflecting selectively impaired membrane targeting and p.H193P reflecting combined expression and trafficking defects. Additionally, NRPE-stimulated ATPase activities were impaired in VUS, p.V195L, p.V195I, p.D197H, p.I214F and p.N269S. Overall structural destabilization interfered with the NRPE-stimulated ATPase activities of p.N269S, while the lack of NRPE-stimulated ATPase activities of p.D197H, p.V195L, p.V195I and p.I214F are thought to be due to impaired NRPE interactions with ABCA4. All the VUS at p.R140, p.H193Y, p.D197N and p.N269H showed both the basal and NRPE-stimulated ATPase activities but less than that of the wild type displaying a mild functional deficit. Together, these findings demonstrated that certain VUS within the unresolved ECD1 region disrupts ABCA4 stability and function, supporting their contribution to disease pathogenesis. This integrative approach highlights key residues likely to be pathogenic and advances the interpretation of VUS in inherited retinal disorders.

## Introduction

Inherited retinal diseases (IRDs) constitute a genetically heterogeneous group of rare disorders characterized by progressive vision loss resulting from abnormal development or degeneration of photoreceptor cells in the retina [1]. The most prevalent forms of IRDs refer to retinitis pigmentosa, cone-rod dystrophy, and Stargardt disease (STGD1) [1–3]. Stargardt disease is the most prominent IRD in children, with clinical and molecular hallmarks, affecting around 1 in 8000 to 10,000 individuals worldwide [4]. It is inherited in an autosomal recessive pattern, marked by early-onset central vision loss with preserved peripheral vision for an extended period in life [3, 4]. STGD1 is clinically characterized by the accumulation of lipofuscin deposits, which appear as pisciform flecks within the retinal pigment epithelium (RPE), which are associated with photoreceptor and RPE cell death [5–7]. The toxic bis-retinoid lipofuscin accumulation is primarily caused by pathogenic variants in the ABCA4 protein in photoreceptors of the retina [8, 9].

The ABCA4 protein, also known as ABCR and the rim protein, is encoded by the *ABCA4* gene (OMIM #601691) located on chromosome 1p22 [8, 10, 11]. ABCA4 is an ATP-powered transporter protein composed of 2273 amino acid residues and is classified within the ATP-binding cassette transporter superfamily [12]. It is expressed in both cone and rod outer segments and is particularly located at the periphery of the photoreceptor disc membranes (Fig 1) [10, 13]. ABCA4 exports N-retinylidene-phosphatidylethanolamine (NRPE), formed from all-trans retinal (ATR) and phosphatidylethanolamine from the disc lumen to the photoreceptor cytosol for their subsequent recycling through the visual cycle in retinal pigment epithelium [12, 14, 15]. The proper transport of retinoid intermediates across the photoreceptor disc membranes is essential to prevent the accumulation of toxic bis-retinoid byproducts and to maintain photoreceptor homeostasis [16].

**Fig 1.**
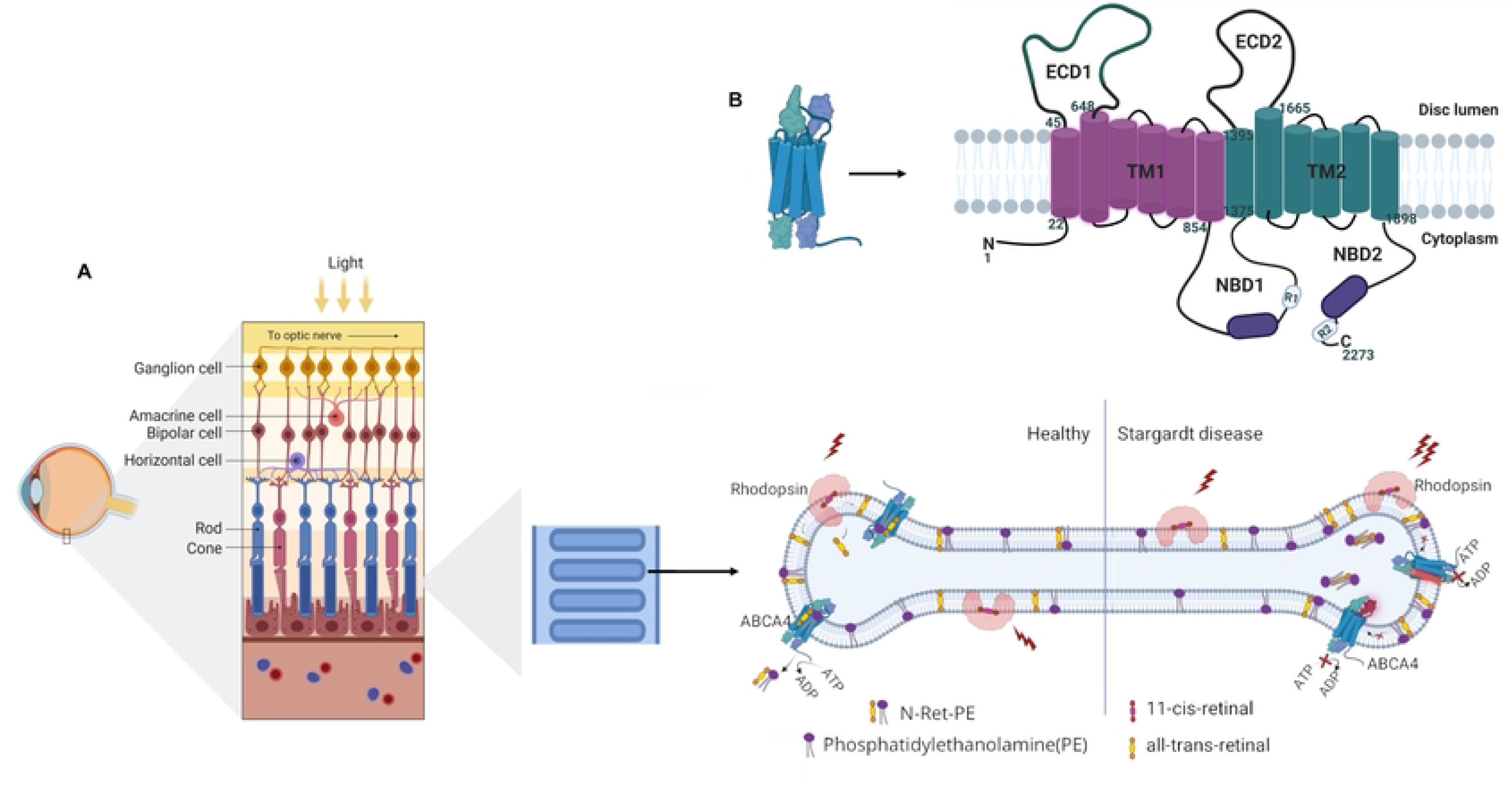
Structural and Functional Role of ABCA4 in Photoreceptor Outer Segment and Stargardt Disease. **A)** Localization of ABCA4 and molecular mechanism of the Stargardt disease. The retina is composed of a layered organization of cell types, including the retinal pigment epithelium (RPE), photoreceptors (cone and rod cells), horizontal cells, bipolar cells, amacrine cells, and ganglion cells. Photoreceptor outer segments contain multiple stacks of discs for efficient phototransduction. ABCA4 is located at the periphery of the disc membranes. 11-cis-retinal bound to Rhodopsin undergoes photoisomerization to all-trans-retinal (ATR) in the presence of light. ATR conjugates with membrane lipid phosphatidylethanolamine (PE) and forms N- retinylidene-phosphatidylethanolamine (NRPE) reversibly. In a healthy retina, NRPE is actively transported from the photoreceptor disc to the cytosol via ABCA4, enabling recycling of 11-cis-retinal in the visual cycle. In Stargardt disease, structural or functional defects in ABCA4 inhibit NRPE transport, leading to accumulation of NRPE in the discs and the buildup of toxic retinoid byproducts, such as lipofuscin. The loss of ATP hydrolysis activity is a characteristic feature of pathogenic ABCA4 variants. **B)** Illustration of the domains and the topology of ABCA4 in photoreceptor disc membranes. ABCA4 exhibits internal pseudo-twofold symmetry with each half containing an extracellular domain (ECD), transmembrane domain (TMD), and a nucleotide-binding domain (NBD). The ECDs are oriented towards the disc lumen, and the NBDs are in the photoreceptor cytosol.

ABCA4 is a highly polymorphic gene that generates a large number of genetic variants in the population, including both benign and pathogenic variants [7, 17]. The pathogenic variants of ABCA4 result in structurally and functionally defective ABCA4 proteins, which impede the transport of retinoid metabolites across the disc membrane [12]. Consequently, NRPE accumulates in the disc lumen, conjugating to form toxic retinoid compounds, such as lipofuscins, thereby increasing retinal oxidative stress and leading to photoreceptor degeneration and central vision loss in STGD1 [6, 18].

Pathogenic variants in the *ABCA4* gene arise from a range of mutation types, including frameshift, nonsense, missense, and splice-site mutations. Among these, null mutations, such as frameshift and nonsense variants, and compound mutations tend to be more deleterious, often resulting in severe, early-onset forms of STGD1 [4, 17, 19]. Given the polymorphic nature of *ABCA4*, missense mutations exhibit a broad phenotypic spectrum, ranging from mild, late-onset disease to severe, early-onset cases that closely resemble those caused by null alleles [4, 20, 21]. This variability complicates genotype–phenotype correlations and presents challenges for clinical interpretation and disease prognosis.

According to the latest count at submission, ClinVar database reports 4634 genetic variants (https://www.ncbi.nlm.nih.gov/clinvar/, accessed June 23, 2026), with missense variants accounting for a significant percentage (∼41% n=1898). Approximately half (51%) of them are categorized as pathogenic/likely pathogenic, or benign/likely benign, while ∼49% (n=931) of missense variants are classified as variants of uncertain significance (VUS) due to lack of or conflicting evidence regarding their pathogenicity [22, 23]. The clinical ambiguity around ABCA4-VUS poses a challenge for patients with such variants participating in clinical trials. Therefore, it is imperative to categorize these variants into more defined categories to facilitate early diagnosis and treatment for patients, as well as aid clinicians in disease diagnosis and management [17, 22].

The major objective of this study is to evaluate the pathogenicity of ABCA4 missense VUS in both computational and experimental platforms. The ABCA4 protein is a single polypeptide exhibiting internal symmetry, composed of two large extracellular domains (ECD1 and ECD2), two transmembrane domains (TMD1 and TMD2), and two nucleotide-binding domains (NBD1 and NBD2) (Fig 2) [10, 13]. Given the larger domain size, the highest proportion (27%) of the VUS are reported in the ECD1 domain, drawing significant attention to this disc lumen-oriented extracytoplasmic domain of ABCA4. ABCA4 extracellular domains (ECD1 and ECD2) together form a lid-tunnel-base configuration in which the lid and the tunnel are formed from ECD1, and the base is formed from ECD2 [13]. Notably, the longest unresolved region in the cryo-EM structures of ABCA4 is located in the apical lid region of ECD1 domain (Fig 2A), spanning approximately residues 138–271 [10, 13]. According to the authors, accurate amino acid assignment and subsequent secondary structure modeling of this segment were limited by the high conformational flexibility of the lumen-exposed lid region of ECD1 (Fig 2A) [10, 13].

**Fig 2.**
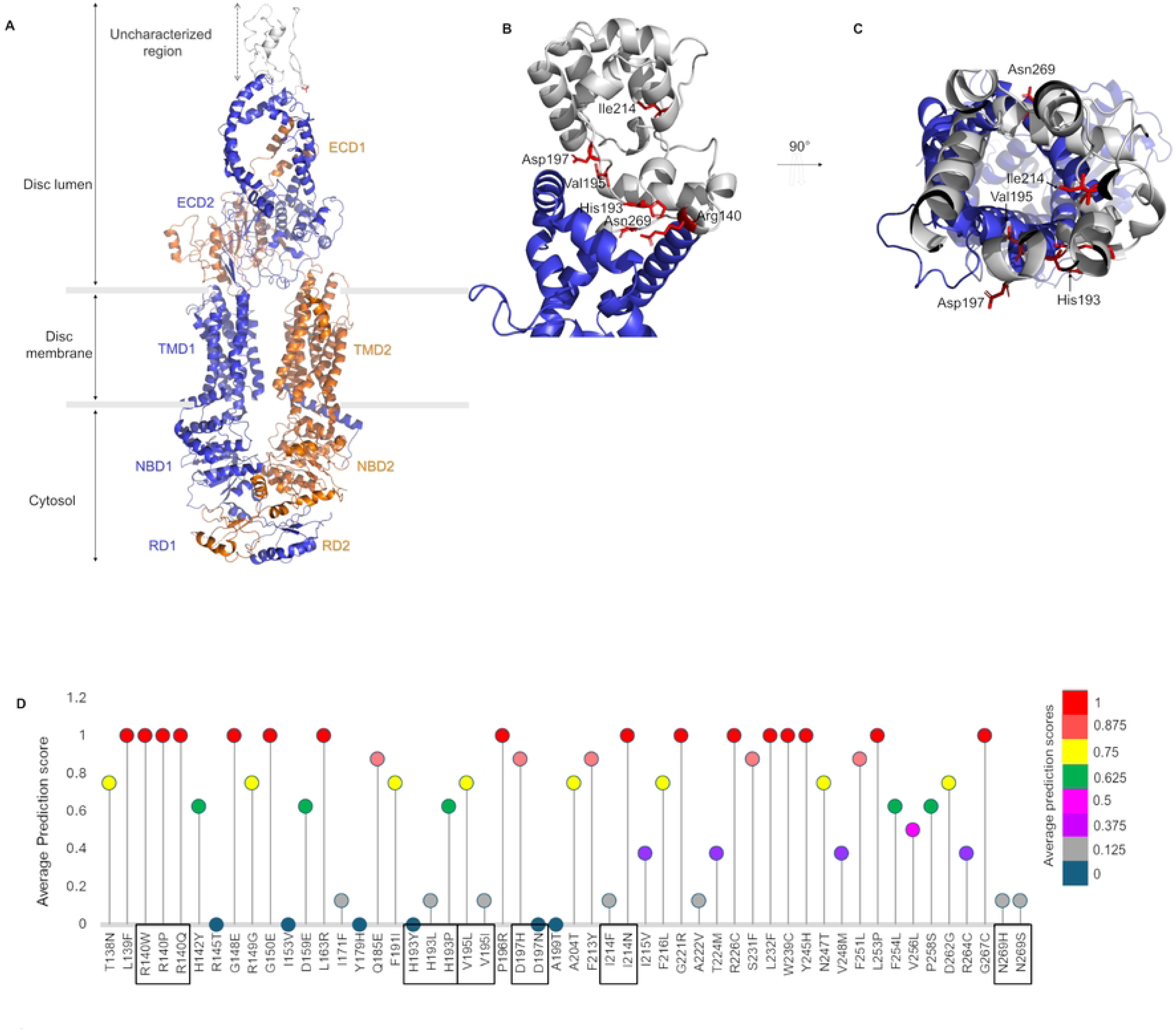
Structural mapping and predicted pathogenicity of ABCA4 missense variants in the uncharacterized region. **A)** The domain organization of ABCA4. The N-terminal and C-terminal halves are shown in blue and orange, respectively. **B)** Localization of the mutation hotspots in the AlphaFold predicted secondary structures of the ECD1 uncharacterized region. **C)** A luminal view of the distribution of mutation hotspots in the AlphaFold predicted secondary structures of the ECD1 uncharacterized region. **D)** A diagram of the average pathogenicity prediction scores for each missense VUS identified in the ECD1 uncharacterized region. Predictions from four *in silico* tools (MetaRNN, MutScore, REVEL, and PolyPhen-2) were used to generate average score representation. Based on the prediction from each tool, scores were assigned: 0-benign/likely benign, 0.5-VUS, and 1-pathogenic/likely pathogenic, and the average is plotted.

The S-loop of ECD1, in close proximity to the NRPE binding site, participates in NRPE binding interactions via its p.R653, p.R587, p.W339, and p.Y345 residues according to prior cryo-EM and functional studies [10, 24]. Additionally, lipid-like molecules were observed in the hydrophobic tunnel of ECDs in ABCA4 cryo-EM studies [10, 13, 24]. Other than that, the specific function or functionally important structural motifs of the elongated tunnel and lid sections of ECD1 are not clearly identified in the literature [10, 24, 25]. Therefore, structural and functional analysis of these unexplored variants in the ECD1 uncharacterized lid region would provide important information for several broad perspectives such as understanding its pathogenic impact towards IRDs, mechanistic studies to unravel its function, ECDs ligand binding interactions and the development of novel therapeutic strategies. We hypothesize that variants in the ECD1 uncharacterized region disrupt ABCA4 function through structural destabilization and impaired NRPE-stimulated ATPase activity. The lack of well-defined secondary structures in the uncharacterized region of ECD1 undermines the reliability of pathogenicity predictions for the VUS in this area using *in silico* tools. Therefore, for robust and comprehensive analysis of the activity and the pathogenicity of these VUS, *in silico* analysis was coupled with experimental platforms, such as functional assays. The characterization of VUS within the uncharacterized region of the ECD1 domain was further focused on a subset of multiallelic sites identified in this region. Each multiallelic site comprises at least two distinct amino acid substitutions classified under the VUS category in ClinVar. All the VUS reported at each multiallelic site were evaluated, enabling controlled comparison of the structural and functional impacts by different side-chain properties at a single locus. In the first stage, the pathogenicity of these variants was assessed using bioinformatic analyses and *in silico* pathogenicity prediction tools. Subsequently, the VUS were expressed in virus-like particles (VLPs) using the baculovirus expression system, and their functional impacts were evaluated using an ATPase activity assay against the ABCA4 wild-type (WT) protein. This study aims not only to establish genotype–phenotype correlations for the identified VUS through functional analyses but also to provide an in-depth evaluation of the distinct pathogenic effects of amino acid substitutions at the multiallelic sites of the ECD1 uncharacterized region and to provide functional evidence for the reclassification of these VUS.

## Materials and methods

### Reagents and buffers

Bacterial growth media 2XYT Broth was from RPI (Cat. No. X15600-10000.0), For bacterial growth plates, agar powder was purchased from Thermo Scientific (Cat. No. A10752-0E) and IPTG was from GOLDBIO (Cat. No. 12481C50). The antibiotics used in bacterial cell culture were purchased from VWR Life Sciences, Kanamycin sulfate (Cat. No. 0408-EU-25G), Gentamycin sulfate (Cat. No. 0304-50G), Ampicillin sodium salt (0339-25G), Tetracycline Hydrochloride (Cat. No. 0422-100G). Cellfectin II reagent for transfection was purchased from Thermofisher (Cat. No. 10362100). Bovine Serum Albumin (BSA) used for western blotting and buffers were from Fisher Bioreagents (Cat. No. BP9706-100). NRPE was prepared using all-*trans* Retinal from Cayman Chemical Company (Cat. No. 18449) and DOPE from Avanti Research (Cat. No. 850725C-25mg). ATP-α-^32^P was purchased from Revvity (Cat. No. BLU003H250UC), the TLC plates were from Sigma-Aldrich (Cat. No. Z122882-25EA), and the scintillation cocktail was from RPI (Cat. No. 111195). Buffers used in the study: Dulbecco’s phosphate-buffered saline (DPBS) from Thermo Fisher Scientific (Cat. No. 14040133); RIPA lysis buffer: 50 mM Tris-HCl (pH 8.0), 150 mM NaCl, 5 mM EDTA, 1% NP-40, 0.5% sodium deoxycholate, 0.1% SDS; Tris-buffered saline (TBS-T): 10 mM Tris-HCl (pH 8.0), 150 mM NaCl and 0.1% Tween 20; NRPE buffer: 25 mM HEPES (pH 8.0), 150 mM NaCl, 5 mM MgCl_2_, 10% Glycerol, 10 mM CHAPS; ATPase assay buffer: 100 mM Tris-HCl (pH 8.0), 500ug/mL BSA, 25 mM DTT, 50% Glycerol.

### AlphaFold 3 protein modeling and variant structural analysis

The wild-type ECD1 domain of ABCA4 was modeled using AlphaFold 3, a deep-learning-based computational framework developed by DeepMind, to predict the secondary structure of the uncharacterized region at the apex of ABCA4 (S1 Fig) [26]. Domain-specific modeling was performed for the ECD1 domain to improve the accuracy and confidence of structural predictions for the uncharacterized region. The ECD1 domain structure with the highest confidence was used for the variant structural analysis.

The WT ECD1 structure was visualized in PyMOL 3.1.3 software (The PyMOL Molecular Graphics System, Version 3.1.3 Schrödinger, LLC, New York, NY, USA). The VUS hotspots were mapped in the ECD1 uncharacterized region, and the secondary structures containing the VUS were identified. The mutagenesis feature in PyMOL was used to generate structures containing each VUS. Once each VUS structure was generated, the neighboring residues located at 4 Å from each mutant residue were visualized, and alterations in intramolecular interactions, such as the loss or formation of hydrogen-bonding interactions and steric clashes with adjacent residues, were identified relative to the respective WT residue. The structural destabilization energies for each VUS were estimated as Gibbs free energy changes (ΔΔG) using the FoldX plug-in in YASARA, followed by energy minimization [27, 28].

### Curation of ABCA4 variants from ClinVar and *in silico* pathogenicity assessment

ClinVar, a publicly available clinical variant database, contained approximately 4,400 reported ABCA4 variants at the time of data retrieval (accessed March 5, 2025). These variants were subsequently filtered to retain missense variants classified as variants of uncertain significance (VUS).The missense VUS reported in the uncharacterized region (138-271) were extracted. There were 52 missense VUS identified (S2 Table), and six loci with more than one missense VUS were identified as multiallelic sites in the uncharacterized region.

The pathogenicity of each of the 52 VUS was evaluated using four high-performing *in silico* prediction algorithms: REVEL, MutScore, MetaRNN, and PolyPhen-2. REVEL, being an ensemble tool, uses 13 prediction algorithms to provide precomputed prediction scores (S2 Table) [29]. PolyPhen-2 uses structural and evolutionary features of proteins and amino acid residues to generate a pathogenicity score. Two novel classifiers, MutScore, known for its high sensitivity, and MetaRNN, specific to rare VUS, were used to enhance prediction precision [30, 31]. The prediction scores of these tools range from 0 to 1, with high scores indicating pathogenicity and lower scores indicating benign variants.

### Insect cell culture

Spodoptera frugiperda 9 (Sf9) (Cat. No. 12659017, Thermo Fisher Scientific) and Trichoplusia ni BTI-TN5B1-4 High Five, (Hi5) (Cat. No. ENH127-FP, Kerafast) cells were cultured in Gibco SF900-III serum-free medium (Thermo Fisher Scientific) without the addition of antibiotics or supplements and maintained at 27 °C in a cell culture incubator.

### ABCA4 variant generation with site-directed mutagenesis

Fourteen ABCA4 missense VUS were generated using site-directed mutagenesis using Quick-change Lightening Site-directed Mutagenesis kit (Agilent, Cat. No. 210519-5) according to the manufacturer’s protocol. pFastBac Dual-ABCA4 (pFBD-ABCA4) vector containing full-length wild-type ABCA4 coding sequence (on the genome assembly GRCh38:Chr1:93, 992, 834-94, 121,148 and Reference Transcript NM_000350.3) was used as the template. The ABCA4 sequence and the mutations in the ABCA4 variants were confirmed by whole-plasmid sequencing.

### Expression and isolation of ABCA4-VLPs using the Baculovirus expression system

Recombinant ABCA4 variant proteins were expressed in VLPs using the Bac-to-Bac baculovirus expression system described in the Invitrogen Life Technologies instruction manual. The first step was to generate a VLP-producing baculovirus vector (bacmid) for each variant. pFBD vector (negative control), pFBD-ABCA4 (WT), and the pFBD-ABCA4 variant vectors were transformed separately into DH10Bac VLP factory cells (Geneva Biotech) to generate the respective bacmids via site-specific transposition of the ABCA4 sequence into the baculovirus shuttle vector in DH10Bac VLP factory *E.coli* cells. Colonies containing the WT/variant sequence were identified by blue-white screening, propagated, and the bacmids were isolated. Polymerase chain reaction was conducted with two ABCA4-specific forward and reverse primers using the isolated bacmids as templates to confirm the successful incorporation of the ABCA4 coding sequence into the baculovirus shuttle vector.

The second step was the generation of VLP-producing baculovirus. In this study, 13 missense VUS located at 6 multiallelic sites were organized into 4 clusters based on their physicochemical properties, enabling appropriate comparisons among variants occurring at the same site. Baculovirus and VLP generation steps were conducted in batches, with each batch containing the negative control (Δ*ABCA4*), ABCA4-WT, and one/two clusters of VUS to achieve better comparison of the variant functional impact. VLP-producing baculovirus was generated by the infection of Sf9 insect cells with bacmid constructs. A batch of bacmids was separately transfected into Sf9 cells in a 6-well plate (1 × 10^6^ cells/well) with 2 µg of bacmid DNA using 6 µL of Cellfectin reagent in SF900-III media, and the cells were incubated at 27 °C for 72 h to produce baculovirus. After 72 h, media containing the initial virus titers (V_0_) were collected by centrifugation (500g, 5 min). Sf9 cells were infected with V_0_ to generate a high-virus titer (V_1_) for VLP production. Confluent (40-60%) Sf9 cells in T75 flasks were infected with 0.5 mL of each V_0_ and incubated at 27 °C for 72 h. After 72 h, V_1_ were collected by centrifugation (500g, 5 min). Quantitative PCR (qPCR) with qPCR primers for ABCA4 was used to quantify the number of ABCA4 copies in each of the V_1_ titers.

The third step was the protein expression in VLPs for negative control, ABCA4-WT, and ABCA4 variants. Adherent Hi5 cells were used for VLP production due to their superior performance in recombinant protein expression. Confluent (40-60%) Hi5 cells in T182 flasks were infected with respective V_1_ s containing 2.5 × 10^8^ copies of ABCA4 and incubated at 27 °C for 2 h with shaking (60 rpm), followed by incubation at 27 °C for 72 h for the production of VLPs. After 72 h, Hi5 cells were harvested, lysed in RIPA buffer, and analyzed by western blot to assess protein expression. The ABCA4-VLPs were isolated from the media by several centrifugation steps. The VLP-containing media were cleared of floating cells by centrifugation at 500g for 10 minutes. Cell debris was removed by centrifugation at 3000g for 30 minutes at 4 °C. The supernatants were then subjected to ultracentrifugation at 30,000 rpm for 4 h at 4 °C to sediment the VLPs. The isolated VLPs were resuspended in 1X PBS and stored on ice at 4 °C.

### Western blot analysis

The expression of the ABCA4 variants in Hi5 cells and their localization to VLPs were detected by western blots, quantified, and the variant-specific impacts on protein expression, folding, and membrane localization were identified compared to ABCA4-WT. Western blots for Hi5 cells and VLPs were developed in batches, each batch containing a negative control, ABCA4-WT, and ABCA4 variants in one/two multiallelic sites. The total protein concentrations of the Hi5 cell lysates and VLPs were determined using the Bradford assay [32].

An amount of 5 µg of total proteins from each sample (cell lysate /VLPs) was resolved on 3-8% tris-acetate precast gels (Invitrogen Life Technologies, Cat. No. EA0378BOX) using SDS-PAGE. Proteins were transferred to a nitrocellulose membrane (Bio-Rad Trans-blot Turbo, Cat. No. 1704158), blocked with 3% BSA in 1X TBS-T for 30 minutes at room temperature. The membranes were incubated with a rabbit anti-ABCA4 primary antibody (Abcam, Cat. No. AB72955) in 1% BSA in 1X TBS-T (1:1000) at room temperature for 1 h. The blot was washed once with 1X TBS-T and incubated with an HRP-conjugated anti-β-tubulin primary antibody (Invitrogen Life Technologies, Cat. No. MA5-16308-HRP) (1:1000) in 1X TBS-T for 1 h at room temperature to detect the loading-control protein (β-tubulin) level in Hi5 cell samples. VLP western blots were incubated with anti-Influenza A M1-Rabbit primary antibody (Invitrogen Life Technologies, Cat. No. PA5-32253) in 1% BSA in 1X TBS-T (1:1000) to detect the loading control protein (Influenza matrix protein) in VLPs. The blots were washed 3 times with 1X TBS-T for 5 minutes each. The blots were incubated with anti-rabbit secondary antibody (Abcam, Cat. No. AB205718) in 1% BSA in 1X TBS-T (1:10,000) for 30 minutes at room temperature, followed by 1X TBST wash. The blots were developed with the Radiance Plus chemiluminescence substrate (Azure Biosystems, Cat. No. AC2103), and the protein bands were detected with the iBright imaging system (Thermo Fisher Scientific). Triplicated western blots were quantified using ImageJ 1.53a software, normalized, and plotted using GraphPad Prism 10.6.1 software.

### Radiolabeled ATPase assay

Retinal byproduct transport across ABCA4 is coupled with ATP hydrolysis. Therefore, the transport function of ABCA4 variants can be measured in terms of their ATPase activities using an established radiolabeled ATPase assay [22, 33]. The ATPase assay was conducted in batches; each batch included VLPs from the negative control, ABCA4-WT, and ABCA4 variants belonging to one/two mutation hotspots for comparison. For each variant, basal ATPase activity without retinal substrate, NRPE, and the retinal-stimulated ATPase activity with NRPE were measured.

NRPE substrate was prepared *in situ* by mixing 0.4 µmol of ATR with 1.2 µmol of DOPE lipid in chloroform. The mixture was incubated at room temperature in the dark for 1 h. The solvent was evaporated under argon, and the product was resuspended in 1 mL of NRPE buffer. An amount of 1 µg of VLP proteins was mixed with 1X TDBG buffer, 5 mM MgCl_2_, 500 µM ATP-α-32P, 40 µM NRPE (retinal stimulation) or NRPE buffer (basal), and the reaction volume was adjusted to 10 µL with distilled water. The reactions were incubated at 37 °C for 1 h in the dark to catalyze ATP hydrolysis. After 1 h, 1 µL of each reaction mixture was spotted on TLC strips pre-spotted with a 5 mM ATP/ADP mixture to enhance visualization of ATP and ADP separation under UV light. The TLCs were developed using a 1 M formic acid and 0.5 M LiCl solution, dried, and visualized under UV light. The ATP and ADP fractions on the TLC plate were excised and added to two separate scintillation vials, each containing 4 mL of scintillation cocktail solution. The radioactivity of the ATP and ADP fractions was measured with a Beckman 6500 liquid scintillation counter. The ATPase activities of all the variants were measured in biological triplicates (n=3). The negative control VLPs contain membrane ATPases other than ABCA4. Therefore, the basal and NRPE-stimulated ATPase activities of the negative control were subtracted from the respective ATPase activities of ABCA4-WT and the ABCA4-variants to obtain the net ATPase activities (pmol/min). The basal and NRPE-stimulated ATPase activities of all the variants, including ABCA4-WT, were normalized based on the ABCA4-WT (basal) ATPase activity in each batch. The % activities were plotted, and statistical analysis was conducted using GraphPad Prism 10.6.1.

### Statistical analysis

Western blot quantifications, basal and retinal-stimulated ATPase activities of the ABCA4-WT, and ABCA4-variants from biological triplicates (n=3) were subjected to statistical analysis using one-way analysis of variance (ANOVA), followed by Tukey’s post-hoc multiple comparison test to identify significant variations in variant expression, membrane localization, and functional activities compared to the ABCA4-WT. The statistical analysis was conducted using the analysis feature in GraphPad Prism 10.6.1. All data are represented as mean ± standard deviation (SD) from at least three independent biological replicates, and adjusted P values are indicated as follows: P < 0.05 (*), P < 0.01 (**), P < 0.001 (***), and P < 0.0001 (****); ns, not significant.

## Results

### AlphaFold 3 based structural prediction for the ECD1 uncharacterized region of ABCA4

AlphaFold is a widely used bioinformatics tool for predicting the 3D structures of proteins from their amino acid sequences. It uses a deep-learning approach to infer the native protein folding patterns [26]. The entire ECD1 domain (from residue 45 to 648) was modeled with AlphaFold 3 to identify possible secondary structures in the uncharacterized region. The precision of the predicted protein structure is measured through several parameters, including per-residue estimate of the confidence (pLDDT) and the predicted template modeling score (pTM). Higher confidence (pLDDT) and modeling scores (pTM) indicate reliable folding performance for the predicted protein structure, which resembles the native protein. The secondary structures of the ECD1 uncharacterized region were predicted with high confidence (pLDDT>90) and a high pTM of 0.75 (S1 Fig). The predicted architecture appears to have five short helices and two intermediate-sized helices connected by long, flexible β-loops (Fig 2 and S1). These alpha helices enclose a hollow cavity, consistent with the ECD1 tunnel [10, 13].

### ABCA4 p.R140 is a structurally critical site in the uncharacterized region of ECD1

ABCA4 p.Arg140 (p.R140) is a multiallelic site located at the beginning of the ECD1 uncharacterized region, specifically on an alpha helix secondary structure according to the predicted AlphaFold structure (Fig 2B). Three different VUS alleles were reported at the p.R140 position in ClinVar: ABCA4 p.Arg140Pro (p.R140P), ABCA4 p.Arg140Trp (p.R140W), and ABCA4 p.Arg140Gln (p.R140Q). This group of mutations explains the structural and functional consequences of the altered positively charged p.R140 side chain into a pyrrolidine loop in p.R140P, an aromatic ring in p.R140W, and a polar aliphatic side chain in p.R140Q. Among these three variants, p.R140P is associated with inborn genetic disease and retinal dystrophy according to ClinVar (Table 1), and one patient was reported with a complex allele bearing p.R140Q {*ABCA4* c. 4667+1 G>T, c.4352+1G>A, c. 419 G>A (p.R140Q)} with damaged retina and the symptom of night blindness [34]. All the *in silico* prediction tools used in this study (MetaRNN, MutScore, REVEL, and PolyPhen-2) predicted these three variants as likely pathogenic (Fig 2D). The agreement in predicted pathogenicity between p.R140 variants across *in silico* prediction platforms strongly suggests that these variants appear to be likely pathogenic. The gold-standard pathogenicity predictor, REVEL, was used to represent pathogenicity scores and predictions for each of the p.R140 variants. According to REVEL, p.R140P was predicted to be likely pathogenic, with the highest REVEL score (0.751) in this group, whereas p.R140W and p.R140Q were predicted to be likely pathogenic, with pathogenic scores of 0.72 and 0.687, respectively. The positively charged side chain of p.R140 interacts with the neighboring tunnel region (Fig 3A) amino acid ABCA4 p. Thr446 (p.T446) via two hydrogen bonds, and one hydrogen bond is observed with the lid region ABCA4 p. Ala192 (p.A192) amino acid (Fig 3A), stabilizing the lid-tunnel interface (Fig 2B) through multiple contacts. The VUS, p.R140P, loses the strong hydrogen-bonding interactions between p.R140 and p.T446 and p.R140 and p.A192, and disrupts the interactions between the lid and the tunnel of the ECD domains (Fig 3A). Additionally, it causes steric clashes with adjacent side chains and may form a kink in the alpha helix it resides in due to the lack of a primary amine in the proline. Therefore, p.R140P causes structural defects in the ECD1 uncharacterized region, leading to a significant structural destabilization (relative ΔΔG 4.78 kcal/mol). This is reflected in the reduced membrane targeting of p.R140P protein to the VLPs, which is significantly lower than in the WT (P=0.0386, Fig 3C). In the second VUS, p.R140W, the tryptophan ring causes severe steric clashes, and loss of hydrogen-bonding interactions with p.T446 and p.A192 as evident in the structural analysis. Despite the structural defects in this variant, the p.R140W protein is expressed in High Five (Hi5) cells and localized to VLPs. In p.R140Q, the polar amide group of the side chain maintains hydrogen bonding interactions with p.A192, which is in close proximity, but loses hydrogen bonding interactions with p.T446 in the tunnel region. The variant p.R140Q exhibits a protein expression pattern similar to that of WT and p.R140W. All three variants showed significantly reduced basal ATPase activities and NRPE-stimulated ATPase activities compared to the WT (P<0.0001, Fig 3D). The reduction of NRPE-stimulated ATPase activities of these variants by ≥ 20% than that of the WT suggest a functional abnormality affecting the efficient substrate transport across them.

**Fig 3.**
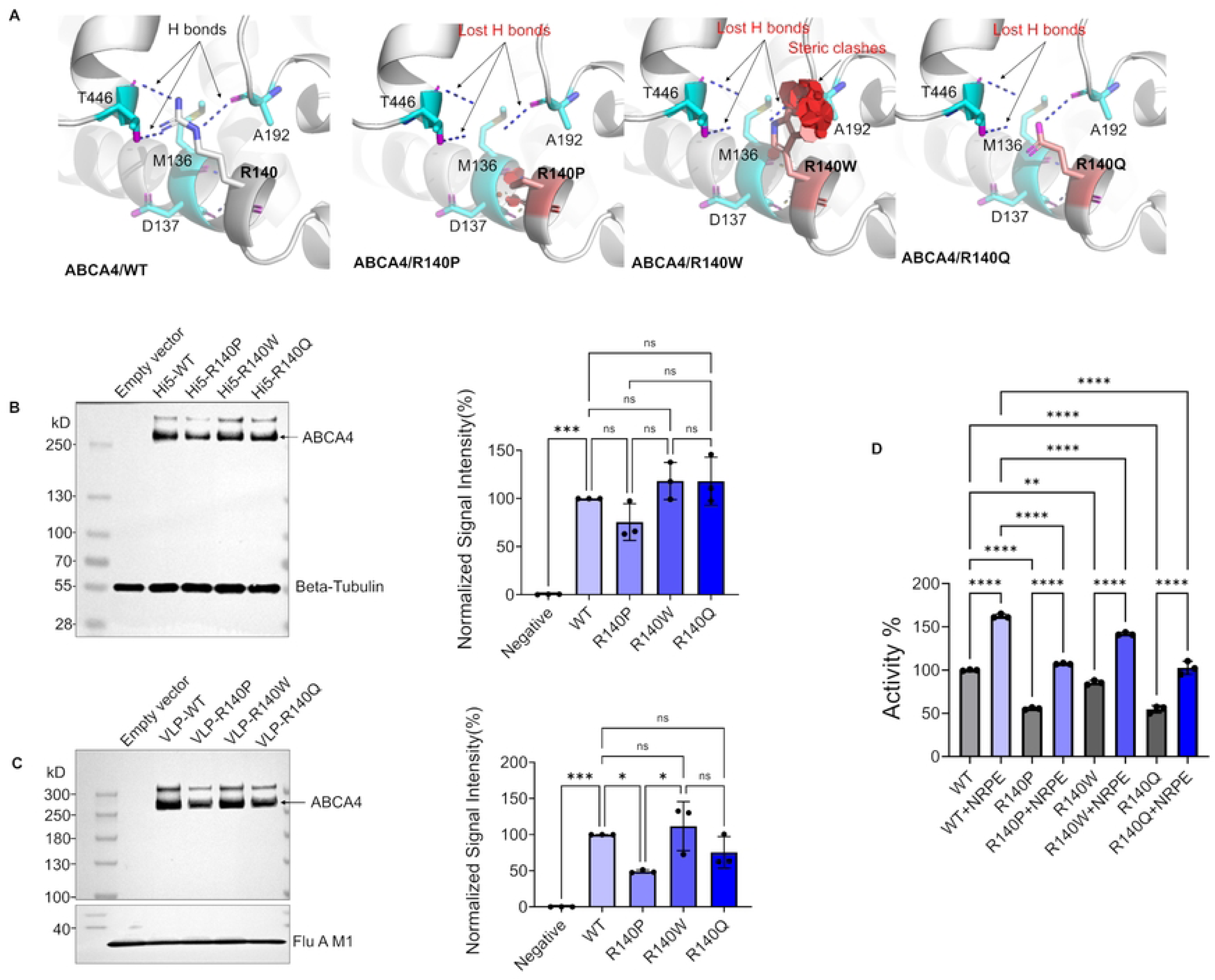
Integrated analysis of ABCA4 p.R140 variants of ABCA4. **A)** Comparative structural analysis of the local interaction environment of the ABCA4 p.R140 residue (a) and p.R140 VUS, generated using the Mutagenesis feature in PyMOL v3.1.3. For each p.R140 VUS, the most probable side-chain rotamer is displayed. The p.R140 side chain is shown in white (carbon) and blue (nitrogen), while neighboring residues are colored cyan (carbon), magenta (oxygen), blue (nitrogen), and yellow (sulfur). Hydrogen-bonding interactions are depicted as blue dashed lines, and steric clashes are indicated by disks (red>yellow>green), with clash severity proportional to the color intensity, size, and number of the disks. WT p.R140; H-bonds with p.M136, p.D137, p.A192, and p.T446, p.R140P; loss of three H-bonds, steric clashes, p.R140W; loss of H-bonds, steric clashes, and p.R140Q; loss of two H-bonds. **B)** Western blot analysis of ABCA4/WT and p.R140 VUS expressed in High Five (Hi5) insect cells, with β-tubulin serving as the loading control. Total protein concentrations of Hi5 cell lysates were determined using the Bradford assay, and equal amounts of protein (5 µg) were loaded per lane. Protein band intensities from biological triplicate experiments (n = 3) were quantified, normalized to β-tubulin, and plotted using GraphPad Prism. **C)** Western blot analysis of ABCA4/WT and ABCA4 p.R140 VUS localized to VLPs, with Influenza A matrix protein 1 (Flu A M1) as the loading control. Total protein concentrations of the VLPs were determined using the Bradford assay, and equal amounts of protein (5 µg) were loaded per lane. Protein band intensities from biological triplicate experiments (n = 3) were quantified, normalized to Flu A M1, and plotted using GraphPad Prism. **D)** Comparative analysis of the basal and NRPE-stimulated ATPase activity of ABCA4/WT and ABCA4 p.R140 VUS VLPs. Total VLP protein (1 µg) from each variant, without NRPE (basal) and with 40 µM NRPE (NRPE-stimulated), was used in the ATPase activity assay. Data are presented as mean ± SD from independent biological replicates (n = 3). Statistical analysis was performed using ordinary one-way analysis of variance (ANOVA), followed by Tukey’s multiple comparisons post hoc test to assess differences between groups. Adjusted P values are indicated as follows: P < 0.05 (*), P < 0.01 (**), P < 0.001 (***), and P < 0.0001 (****); ns, not significant.

**Table 1.**
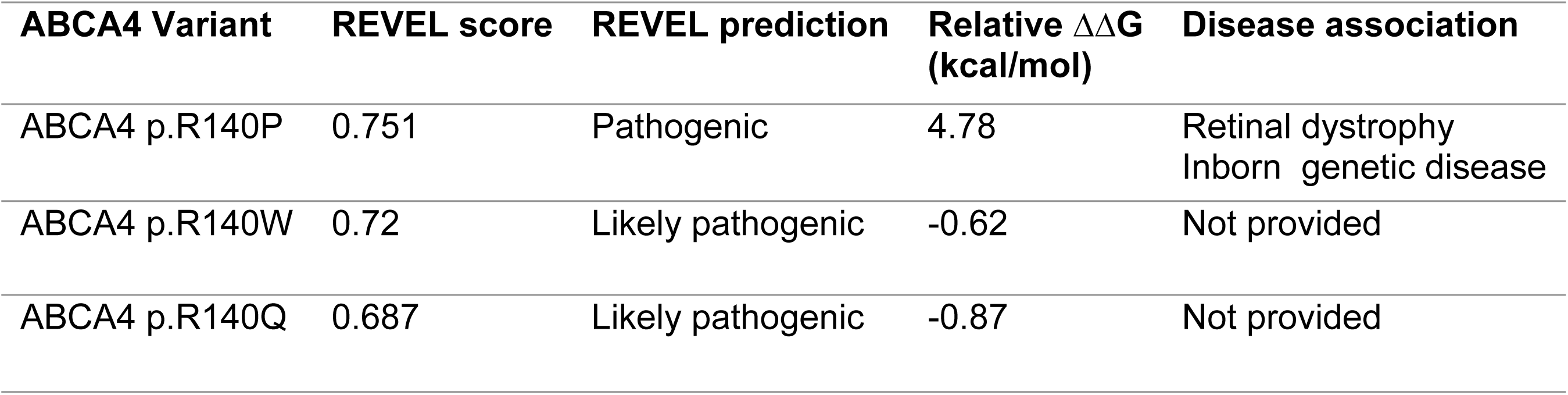
*In Silico* analysis and disease association of ABCA4 p.R140 variants.

### ABCA4 p.H193 site impacts the membrane targeting of the ABCA4 protein

ABCA4 p.His193 (p.H193) is a multiallelic site in the ECD1 uncharacterized region with a polar, positively charged aromatic amino acid in the WT. Residue p.H193 also occurs at the lid-tunnel interface of ECD1, specifically on a loop structure as predicted by AlphaFold 3. The side chain of p.H193 is a solvent-exposed residue and does not make any polar interactions with adjacent amino acid residues (Fig 2C). Therefore, *in silico* and in vitro analyses of the p.H193 VUS provide important insights into the functional role of the ECD1/ECD1 uncharacterized region. This p.H193 locus was reported with three VUS in ClinVar as ABCA4 p.His193Tyr (p.H193Y), ABCA4 p.His193Pro (p.H193P), and ABCA4 p.His193Leu (p.H193L), with no distinct disease association to retinal dystrophies. On the *in silico* platform, p.H193Y was predicted as likely benign by all four prediction tools, and p.H193L was also predicted as likely benign by all but REVEL, which placed it in the VUS score range (Table 2). The only variant predicted to be likely pathogenic by REVEL and MetaRNN in this group is p.H193P (Table 2), suggesting a disease association.

**Table 2.** *In Silico* analysis and disease association of ABCA4 p.His193 variants.

| ABCA4 Variant | REVEL score | REVEL Prediction | Relative $\Delta\Delta G$ (kcal/mol) | Disease association |
| --- | --- | --- | --- | --- |
| ABCA4 p.H193Y | 0.239 | Likely benign | -0.21 | Not provided |
| ABCA4 p.H193P | 0.536 | Likely pathogenic | 3.62 | Not provided |
| ABCA4 p.H193L | 0.442 | VUS | -0.08 | Not provided |

The VUS p.H193Y showed a significant increase in NRPE-stimulated ATPase activity compared to the basal ATPase activity, similar to WT (P<0.0001, Fig 4D). The NRPE stimulation of the p.H193Y against the basal ATPase activity appeared to be maintained by the preserved aromaticity of the variant side chain. But the NRPE-stimulated ATPase activity of p.H193Y is reduced by greater than 20% that of the WT due to the reduced protein expression and the localization to the VLPs. The VUS, p.H193P, showed basal and NRPE-stimulated ATPase activities similar to the negative control VLPs (VLPs prepared from empty bacmid), resulting in a loss in net basal and NRPE-stimulated ATPase activities (Fig 4D). This could be due to the dramatic reduction in p.H193P protein levels in the VLPs (Fig 4C). Additionally, the VUS p.H193P showed several pathogenic traits such as high destabilization energy (relative ΔΔG 3.62 kcal/mol) associated with steric clashes and altered loop structure and reduced membrane targeting. Downstream evaluation of p.H193L VUS was excluded due to its undetectable virus titers. Poor membrane localization was common for the VUS evaluated at p.H193 suggesting its essential role in membrane targeting of ABCA4 in the photoreceptors.

**Fig 4.**
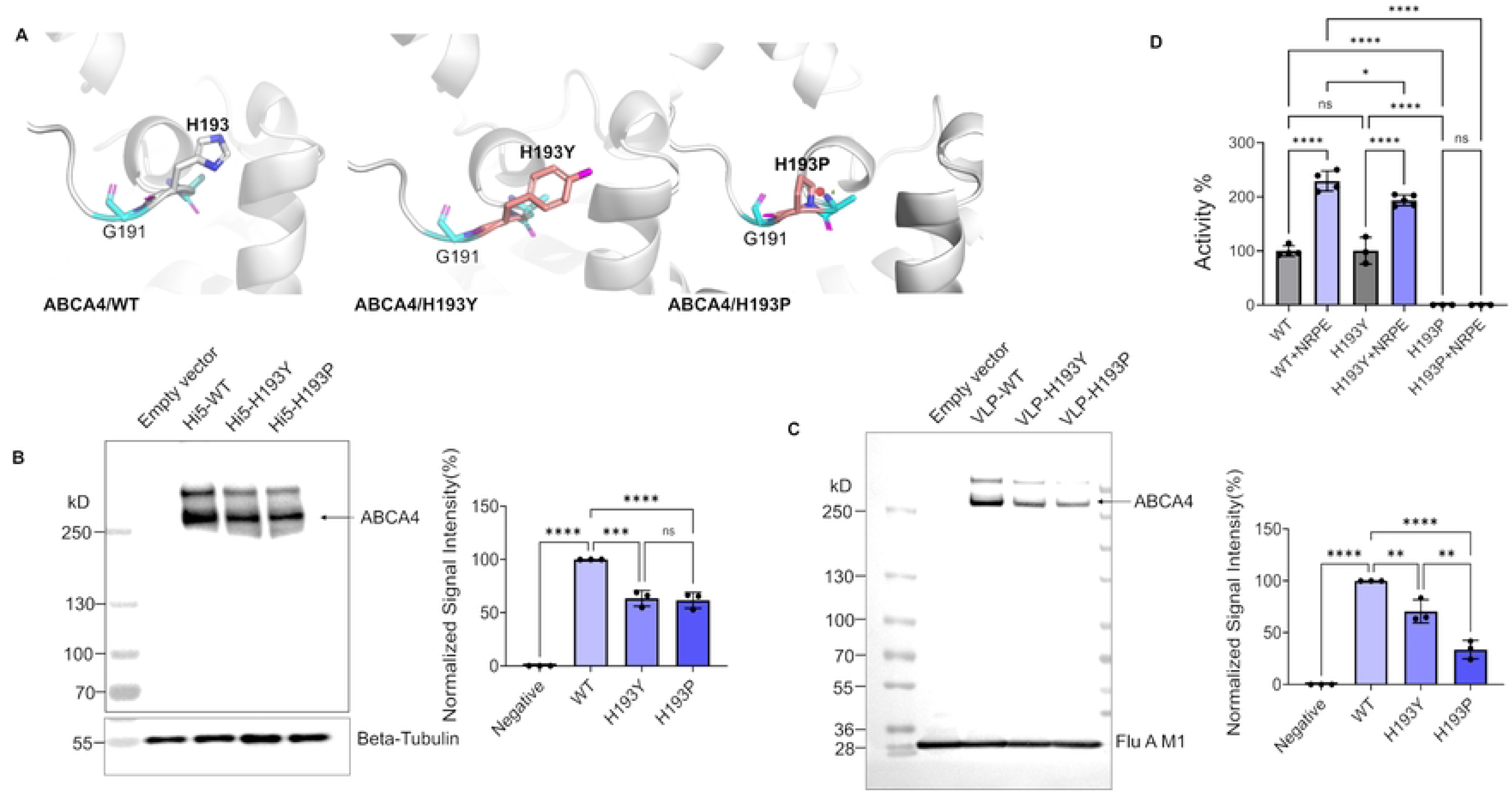
Integrated analysis of ABCA4 p.H193 variants of ABCA4. **A)** Comparative structural analysis of the local interaction environment of the ABCA4 p.H193 residue and p.H193 VUS, generated using the Mutagenesis feature in PyMOL v3.1.3. For each p.H193 VUS, the most probable side-chain rotamer is displayed. The p.H193 side chain is shown in white (carbon) and blue (nitrogen), while neighboring residues are colored cyan (carbon), magenta (oxygen), blue (nitrogen), and yellow (sulfur). Steric clashes are indicated by disks (red>yellow>green), with clash severity proportional to the color intensity, size, and number of the disks. WT p.H193; solvent exposed residue, p.H193Y; no steric clashes, p.H193P; steric clashes, and p.H193L; no steric clashes. **B)** Western blot analysis of ABCA4/WT and p.H193 VUS expressed in High Five (Hi5) insect cells, with β-tubulin serving as the loading control. Total protein concentrations of Hi5 cell lysates were determined using the Bradford assay, and equal amounts of protein (5 µg) were loaded per lane. Protein band intensities from biological triplicate experiments (n = 3) were quantified, normalized to β-tubulin, and plotted using GraphPad Prism. **C)** Western blot analysis of ABCA4/WT and ABCA4 p.H193 VUS localized to VLPs, with Influenza A matrix protein 1 (Flu A M1) as the loading control. Total protein concentrations of the VLPs were determined using the Bradford assay, and equal amounts of protein (5 µg) were loaded per lane. Protein band intensities from biological triplicate experiments (n = 3) were quantified, normalized to Flu A M1, and plotted using GraphPad Prism. **D)** Comparative analysis of the basal and NRPE-stimulated ATPase activity of ABCA4/WT and ABCA4 p.H193 VUS VLPs. Total VLP protein (1 µg) from each variant, without NRPE (basal) and with 40 µM NRPE (NRPE-stimulated), was used in the ATPase activity assay. Data are presented as mean ± SD from independent biological replicates (n = 3). Statistical analysis was performed using ordinary one-way analysis of variance (ANOVA), followed by Tukey’s multiple comparisons post hoc test to assess differences between groups. Adjusted P values are indicated as follows: P < 0.05 (*), P < 0.01 (**), P < 0.001 (***), and P < 0.0001 (****); ns, not significant.

### ABCA4 p.D197 and ABCA4 p.N269 sites are important to maintain the structural stability of ABCA4

The multiallelic sites ABCA4 p.Asp197 (p.D197) and ABCA4 p.Asn269 (p.N269) were studied together because both are polar aliphatic amino acids located in loop regions of the ECD1 uncharacterized region. The residue p.D197 is a negatively charged, solvent-exposed residue according to the AlphaFold structure, which can interact with ions, water, and other proteins and does not interact with the adjacent ABCA4 amino acid residues (Fig 2C and 5A). Two missense VUS, ABCA4 p.Asp197His (p.D197H) and ABCA4 p.Asp197Asn (p.D197N), were reported in ClinVar, with no association with retinal dystrophy. On the *in silico* platform, p.D197H was predicted to be likely pathogenic by three prediction tools, and p.D197N was predicted to be benign by all tools (Fig 2D), as reflected in the REVEL scores and REVEL predictions (Table 3).

**Fig 5.**
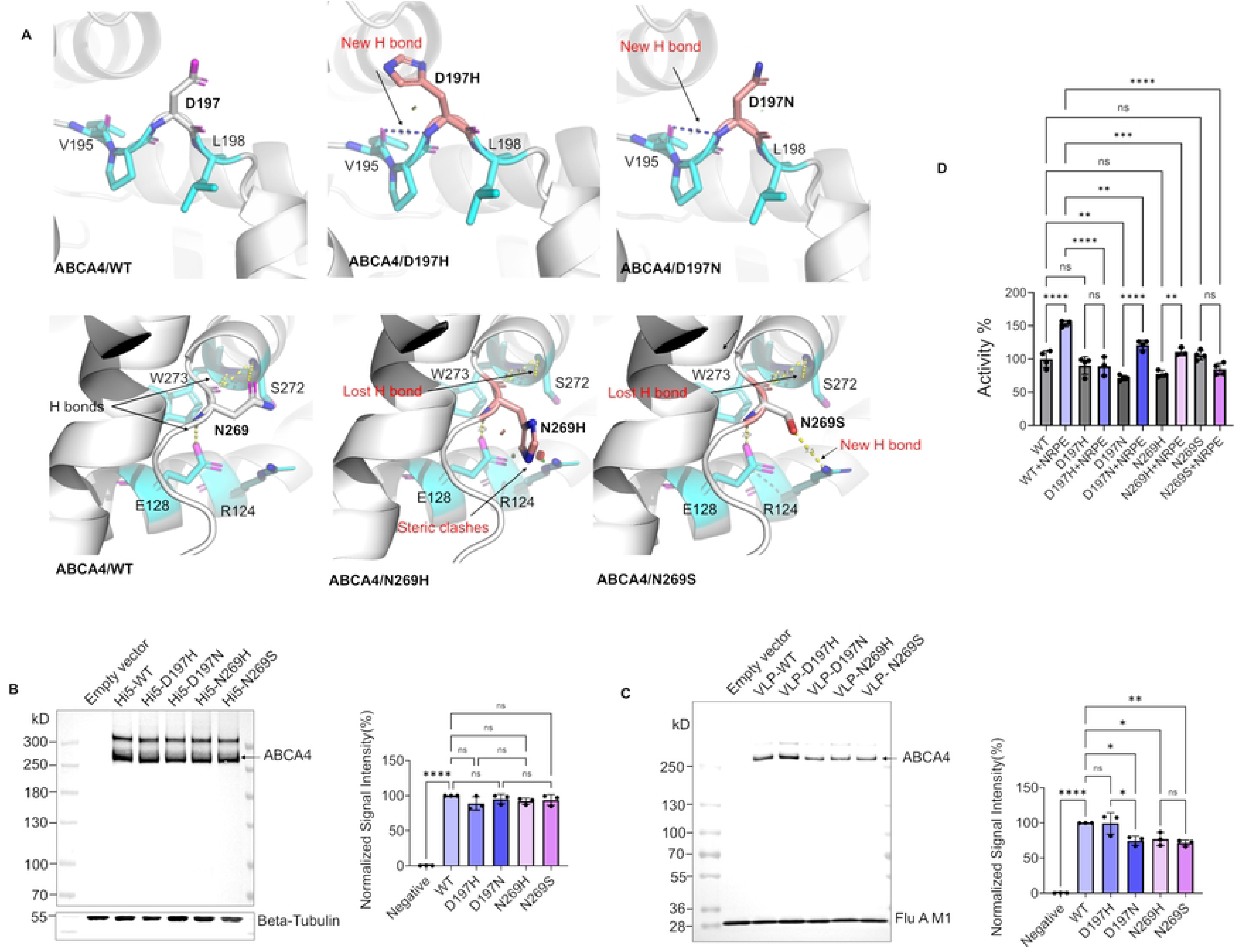
Integrated analysis of ABCA4 p.D197 and ABCA4 p.N269 variants of ABCA4. **A)** Comparative structural analysis of the local interaction environment of the ABCA4 p.D197 and ABCA4 p.D197 VUS, ABCA4 p.N269 and ABCA4 p.N269 VUS, generated using the Mutagenesis feature in PyMOL v3.1.3. For each p.D197 and p.N269 VUS, the most probable side-chain rotamer is displayed. The WT p.D197 and WT p.N269 side chains are shown in white (carbon) and blue (nitrogen), while neighboring residues are colored cyan (carbon), magenta (oxygen), blue (nitrogen), and yellow (sulfur). Hydrogen bonds are shown in blue dashed lines, and steric clashes are indicated by disks (red>yellow>green), with clash severity proportional to the color intensity, size, and number of the disks. WT p.D197; solvent exposed residue, p.D197H; a new H-bond and steric clashes, p.D197N; a new H-bond, WT p.N269; H-bonds with p.E128, p.S272, and p.W273, p.N269H; loss of one H-bond with p.S272 and steric clashes, p.N269S; loss of one H-bond with p.S272 and a new H-bond with p.R124 **B)** Western blot analysis of ABCA4/WT, p.D197 and p.N269 VUS expressed in High Five (Hi5) insect cells, with β-tubulin serving as the loading control. Total protein concentrations of Hi5 cell lysates were determined using the Bradford assay, and equal amounts of protein (5 µg) were loaded per lane. Protein band intensities from biological triplicate experiments (n = 3) were quantified, normalized to β-tubulin, and plotted using GraphPad Prism. **C)** Western blot analysis of ABCA4/WT, p.D197, and p.N269 VUSs localized to VLPs, with Influenza A matrix protein 1 (Flu A M1) as the loading control. Total protein concentrations of the VLPs were determined using the Bradford assay, and equal amounts of protein (5 µg) were loaded per lane. Protein band intensities from biological triplicate experiments (n = 3) were quantified, normalized to Flu A M1, and plotted using GraphPad Prism. **D)** Comparative analysis of the basal and NRPE-stimulated ATPase activity of ABCA4/WT and ABCA4 p.D197 and ABCA4 p.N269 VUS VLPs. Total VLP protein (1 µg) from each variant, without NRPE (basal) and with 40 µM NRPE (NRPE-stimulated), was used in the ATPase activity assay. Data are presented as mean ± SD from independent biological replicates (n = 3). Statistical analysis was performed using ordinary one-way analysis of variance (ANOVA), followed by Tukey’s multiple comparisons post hoc test to assess differences between groups. Adjusted P values are indicated as follows: P < 0.05 (*), P < 0.01 (**), P < 0.001 (***), and P < 0.0001 (****); ns, not significant.

**Table 3.** *In Silico* analysis and disease association of ABCA4 p.Asp197 and ABCA4 p.Asn269 variants.

| ABCA4 Variant | REVEL score | REVEL prediction | Relative $\Delta\Delta G$ (kcal/mol) | Disease Association |
| --- | --- | --- | --- | --- |
| ABCA4 p.D197H | 0.695 | Likely pathogenic | 0.55 | Not provided |
| ABCA4 p.D197N | 0.249 | Likely benign | 0.44 | Not provided |
| ABCA4 p.N269H | 0.495 | VUS | 0.29 | Inborn genetic disease |
| ABCA4 p.N269S | 0.363 | Likely benign | 1.24 | Not provided |

The VUS p.D197H replaces the acidic aliphatic side chain of aspartic acid with the basic aromatic side chain of histidine, shifting the net charge from negative to positive and causing mild steric clashes with the bulky aromatic ring. Furthermore, a new hydrogen bond is formed between the peptide backbone of p.D197H and adjacent p.V195. The structural destabilization caused by these perturbations is reflected in the relative ΔΔG of 0.55 kcal/mol (Table 3). The VUS p.D197N introduces a polar, uncharged aliphatic side chain, altering the negative charge of p.D197 in WT. Despite the polarity change, this is a structurally conserved mutation, as the side chains are similar in size and no steric clashes are observed. But it might affect electrostatic interactions with hydrophilic compounds, such as ions, as well as interactions with other proteins. The variant protein expressions for p.D197H and p.D197N in Hi5 cells remained similar to WT (Fig 5B). However, the p.D197N protein level was lower than that of p.D197H in the VLPs (P=0.0263, Fig 5C). The basal ATPase activities of p.D197H and p.D197N were statistically equal to the basal ATPase activity of WT. But p.D197H failed to show NRPE-stimulated ATPase activity, suggesting an NRPE interaction/transport defect with p.D197H, a likely pathogenic feature, whereas p.D197N showed significant stimulation of ATPase activity (P<0.0001) in the presence of NRPE. The NRPE-stimulated ATPase activity of p.D197N is less than that of the WT, which can be attributed to the lowered VLP targeting of the variant (Fig 5C).

The residue ABCA4 p.Asn269 (p.N269) is located towards the end of the uncharacterized region. It interacts with the ABCA4 p.Glu128 (p.E128) side chain and the ABCA4 p.Ser272 (p.S272) and ABCA4 p.Trp273 (p.W273) backbones of the tunnel region via hydrogen bonds. The multiallelic site p.N269 was associated with two missense VUS, ABCA4 p.Asn269His (p.N269H) and ABCA4 p.Asn269Ser (p.N269S). Both variants were predicted as benign by three of the *in silico* prediction tools and as VUS by one of the four tools used in this study (Fig 2D). REVEL classifies p.N269H as a VUS and p.N269S as likely benign for inherited retinal diseases. On the contrary, p.N269H was reported in a case of an inborn genetic disease in ClinVar, and the p.N269S variant has a high relative destabilization energy of 1.24 kcal/mol (Table 3).

In p.N269H, the polar aliphatic side chain in WT is replaced by a positively charged, bulky aromatic ring, introducing steric clashes and resulting in the loss of one hydrogen-bond interaction between p.N269H and p.S272 in the structural analysis. These structural effects cause mild structural destabilization, as shown by a ΔΔG of 0.29 kcal/mol (Table 3). Although the variant expression in Hi5 insect cells was similar to WT, its VLP targeting is reduced (P=0.0412) (Fig 5B and 5C). The VUS p.N269H showed significantly increased NRPE-stimulated ATPase activity compared to the basal activity (P=0.0048), suggesting that its NRPE stimulation is maintained through mutation. However, the NRPE-stimulated ATPase activity of p.N269H is reduced significantly than that of the WT (P=0.0002) due to lower VLP targeting of p.N269H (Fig 5C).

When p.N269 is mutated to a serine in p.N269S VUS, one of the native hydrogen bonds observed between p.N269 and p.S272 is lost, and a new hydrogen bond is formed with the adjacent p.R124 side chain, leading to a significant structural change resulting in a destabilized conformation evident by the high destabilization energy observed for p.N269S (Table 3). Although the variant expression was similar to WT, the membrane localization was significantly affected (P=0.0094, Fig 5C). Furthermore, the variant failed to stimulate ATPase activity in the presence of the NRPE substrate compared to the basal activity (Fig 5D). The notable destabilizing conformational changes associated with p.N269S might interfere with the NRPE transport mechanisms across ABCA4. The VUS reported on p.D197 and p.N269 caused overall structural destabilizations leading to impaired NRPE-stimulated ATPase activities via affected membrane targeting and NRPE transport mechanisms.

### ABCA4 p.V195 and ABCA4 p.I214 are critical sites associated with NRPE-stimulated ATPase activity of ABCA4

There were two non-polar aliphatic multiallelic sites in the ECD1 uncharacterized region of ABCA4. One of the sites, ABCA4 p.Val195 (p.V195), is located at the interface between the lid and the tunnel of ECD1, along with other polar multiallelic sites on a loop structure (Fig 2B). Two missense VUS were reported at this location in ClinVar: ABCA4 p.Val195Leu (p.V195L) and ABCA4 p.Val195Ile (p.V195I). The p.V195L variant was predicted to be likely pathogenic by REVEL, MetaRNN, and PolyPhen-2, and the average pathogenicity of p.V195L weighs towards likely pathogenic prediction. The VUS p.V195I was predicted as a VUS by REVEL and as benign by other *in silico* tools (Table 4). Therefore, the average prediction for p.V195I favors likely benign nature (Fig 2D). Nevertheless, both of these variants are associated with retinal dystrophy according to ClinVar (Table 4). In both VUS, the side-chain polarity of the site remains unchanged, as both mutations are non-polar, similar to valine, and the only structural variation observed is an increase in the number of carbon atoms and the branching pattern of the carbon chain. These variations do not cause any structural clashes or destabilization, as indicated by the relative ΔΔG values (Table 4).

**Table 4.** *In Silico* analysis and disease association of ABCA4 p.Val195 and ABCA4 p.Ile214 variants.

| ABCA4 Variant | REVEL scores | REVEL prediction | Relative $\Delta\Delta G$ (kcal/mol) | Disease Association |
| --- | --- | --- | --- | --- |
| ABCA4 p.V195L | 0.636 | Likely pathogenic | -0.61 | Retinal dystrophy |
| ABCA4 p.V195I | 0.418 | VUS | -0.4 | Retinal dystrophy |
| ABCA4 p.I214F | 0.375 | Likely benign | -0.04 | Inborn genetic disease |
| ABCA4 p.I214N | 0.683 | Likely pathogenic | 2.72 | Not provided |

The variant expression levels of both p.V195L and p.V195I in Hi5 insect cells are equal to each other, and p.V195I expression is reduced than the WT (P=0.0153, Fig 6B). The p.V195L and p.V195I variant protein levels in the VLPs are equal and do not differ significantly from the WT level in the VLPs (Fig 6C). The VLP targeting of these variants suggests that they follow the membrane localization pattern of WT. The ATPase activities of p.V195L were significantly reduced compared to the WT (basal P=0.0001, NRPE-stimulated P<0.0001). Both p.V195L and p.V195I were unable to demonstrate NRPE-stimulated ATPase activities, suggesting a likely pathogenic mechanism associated with NRPE binding (Fig 6D). Despite structural stability, and membrane localization, these p.V195 variants were functionally defective, consistent with their association with disease. Since a direct NRPE-binding interaction at this p.V195 site has not been identified, the VUS at this site appears to allosterically regulate NRPE binding to ABCA4.

**Fig 6.**
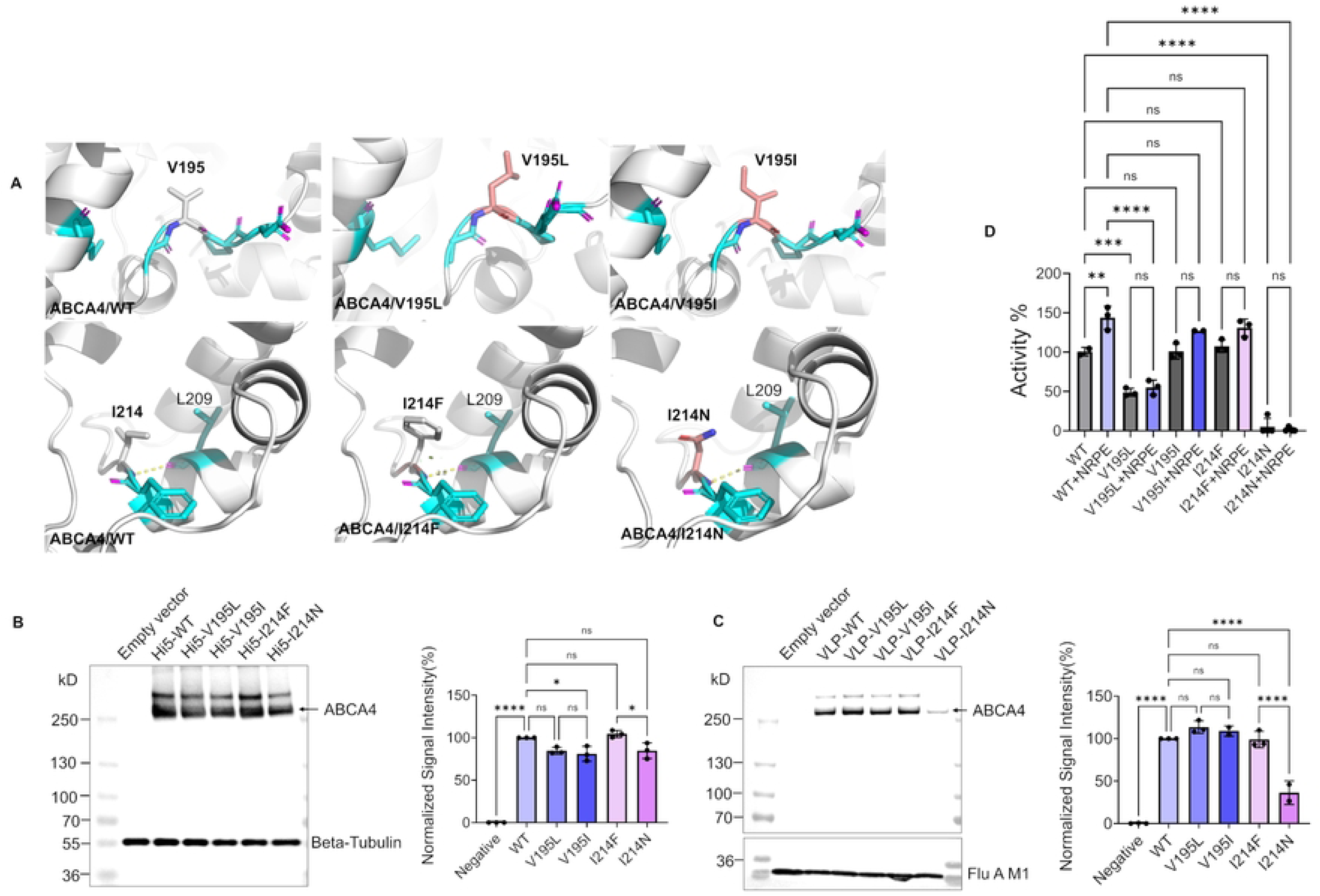
Integrated analysis of ABCA4 p.V195 and ABCA4 p.I214 variants of ABCA4. **A)** Comparative structural analysis of the local interaction environment of the ABCA4 p.V195 and ABCA4 p.V195 VUS, ABCA4 p.I214 and ABCA4 p.I214 VUS generated using the Mutagenesis feature in PyMOL v3.1.3. For each p.V195 and p.I214 VUS, the most probable side-chain rotamer is displayed. The WT p.V195 and WT p.I214 side chains are shown in white (carbon) and blue (nitrogen), while neighboring residues are colored cyan (carbon), magenta (oxygen), blue (nitrogen), and yellow (sulfur). Hydrogen bonds are shown in yellow dashed lines, and steric clashes are indicated by colored disks (red>yellow>green), with clash severity proportional to the color intensity, size, and number of the disks. WT p.V195, p.V195L, and p.V195I variants lack any H-bonding interactions. WT p.I214, p.I214F, and p.I214N show one H-bond between their α-amine nitrogen and the α-carbonyl oxygen of the adjacent p.L209 residue. p.I214F shows mild steric clashes with p.F213. **B)** Western blot analysis of ABCA4/WT, p.V195, and p.I214 VUS expressed in High Five (Hi5) insect cells, with β-tubulin serving as the loading control. Total protein concentrations of Hi5 cell lysates were determined using the Bradford assay, and equal amounts of protein (5 µg) were loaded per lane. Protein band intensities from biological triplicate experiments (n = 3) were quantified, normalized to β-tubulin, and plotted using GraphPad Prism. **C)** Western blot analysis of ABCA4/WT, p.V195, and p.I214 VUS localized to VLPs, with Influenza A matrix protein 1 (Flu A M1) as the loading control. Total protein concentrations of the VLPs were determined using the Bradford assay, and equal amounts of protein (5 µg) were loaded per lane. Protein band intensities from biological triplicate experiments (n = 3) were quantified, normalized to Flu A M1, and plotted using GraphPad Prism. **D)** Comparative analysis of the basal and NRPE-stimulated ATPase activity of ABCA4/WT and ABCA4 p.V195 VUS and ABCA4 p.I214 VLPs. Total VLP protein (1 µg) from each variant, without NRPE (basal) and with 40 µM NRPE (NRPE-stimulated), was used in the ATPase activity assay. Data are presented as mean ± SD from independent biological replicates (n = 3). Statistical analysis was performed using ordinary one-way analysis of variance (ANOVA), followed by Tukey’s multiple comparisons post hoc test to assess differences between groups. Adjusted P values are indicated as follows: P < 0.05 (*), P < 0.01 (**), P < 0.001 (***), and P < 0.0001 (****); ns, not significant.

The other non-polar VUS multiallelic site was ABCA4 p.I214 (p.I214). It is located in the top half of the ECD1 uncharacterized lid region on a loop structure. The non-polar side chain orients to the inner cavity which is covered by several short helices (Fig 2B and 2C). The p.I214 residue of WT interacts with the adjacent ABCA4 p.Leu209 (p.L209) via a hydrogen bond between their backbone atoms. Two missense VUS, p.Ile214Phe (p.Ile214F) and ABCA4 p.Ile214Asn (p.I214N), were reported in ClinVar for this non-polar site. The VUS p.I214F was predicted to be likely benign (0.375) by REVEL, in agreement with other *in silico* prediction tools (Fig 2D), and was associated with an inborn genetic disease. The VUS p.I214N was predicted as likely pathogenic in REVEL (0.683) and in all other prediction tools (Fig 2D) with no reported disease association. In the VUS p.I214F, a mild steric clash is observed with the neighboring ABCA4 p.Phe213 (p.F213) due to the increased steric bulk of the aromatic rings. With p.I214N, the side-chain polarity changes from nonpolar to polar, resulting in a significant alteration. Although no novel polar interactions were observed with p.I214N, it causes an overall structural destabilization, with relative ΔΔG of 2.72 kcal/mol (Table 4), suggesting a pathogenic setting for p.I214N.

In Hi5 insect cells, both p.I214F and p.I214N variants express similarly to WT, but p.I214N expression is reduced compared to p.I214F (P=0.0114, Fig 6B). Furthermore, the p.I214N level in VLPs is dramatically reduced compared with WT and p.I214F (P<0.0001), indicating that it is not properly localized to VLPs (Fig 6C). NRPE-stimulated ATPase activity of p.I214F was not significant compared to the basal activity, suggesting that it does not confer functional activity in NRPE transfer across the variant. Additionally, p.I214N ATPase activities were similar to those of the negative control VLPs, yielding no net basal or NRPE-stimulated ATPase activities due to its poor targeting potential to the VLPs. The functional inactivation of p.I214N is attributed to its conformational destabilization and the altered side-chain polarity. The disease association of p.I214F and the *in silico* pathogenicity prediction for p.I214N align with the functional defects observed in these two variants.

## Discussion

ABCA4 has several uncharacterized regions spanning the ECDs, TMDs, and regulatory domains (RDs), and the major uncharacterized region is located towards the N-terminus of ECD1, as evidenced by cryo-EM structures [10, 13]. An integrated *in silico* and in vitro analysis approach was used to characterize missense VUS reported in the uncharacterized region of ECD1 to unravel their pathogenic potential in IRDs. Among the missense VUS sites in this uncharacterized region, 6 multiallelic sites were identified: p.R140, p.H193, p.V195, p.D197, p.I214, and p.N269. This study characterized 13 missense VUS found at the 6 multiallelic sites in the ECD1 uncharacterized region. The VUS at multiallelic sites were selected to systematically assess how distinct amino acid substitutions at a single locus differentially influence local structural stability and ABCA4 functional activity based on their individual physicochemical properties. Fourteen VUS span across the 6 multiallelic sites were generated; however, 13 VUS were analyzed with p.H193L excluded from downstream analyses due to an undetectable baculovirus titer during expression. Nevertheless, additional variants reported post-study at the same multiallelic sites were not captured in the present analysis.

The secondary structures of the ECD1 uncharacterized region were predicted with high confidence and modeling scores from AlphaFold 3. According to the predicted structural arrangement, all these multiallelic sites are located at the lid-tunnel interface of ECD1, except p.I214, which is located at the top of the lid region. The sites p.R140 and p.N269 mark the N- and C-terminal ends of the uncharacterized region (Fig 2B) and interact with residues from both the lid and tunnel regions (Fig 3A and 5A), serving as two connectors between the lid and tunnel of ECD1. However, the structural confidence metrics generated by AlphaFold are based on internal model assessments rather than direct experimental validation. Cryo-EM data indicate that this uncharacterized region is highly flexible; thus, the static conformation predicted by AlphaFold may not accurately capture the dynamic nature of the ECD1 lid region in the native protein. The structural destabilization energies (ΔΔG) for the VUS were also estimated using the AlphaFold structure in FoldX. Due to the reduced reliability of the estimated ΔΔG values in structurally flexible regions, these measurements were reported as approximate or relative destabilization energies. Molecular dynamics and molecular docking studies would provide additional information about the structural organization and ligand-binding interactions in this region. Furthermore, identification of the posttranslational modifications in this uncharacterized region would help elucidate structurally and functionally important motifs.

*In Silico* pathogenicity predictions for the ABCA4 variants provide evidence supporting pathogenicity under PP3 criteria and benignity under BP4 criteria for variant classification according to ACMG standards and guidelines [35]. To qualify as strong supporting evidence, variant pathogenicity or benignity should be consistently predicted by multiple computational algorithms that incorporate diverse features, including evolutionary conservation, physicochemical properties, and sequence context. Therefore, four *in silico* tools (MetaRNN, MutScore, REVEL, and Polyphen-2) with different inputs were used, in accordance with ACMG guidelines. Among them, REVEL was highlighted because it is a widely validated, high-performing ensemble predictor of rare missense variant pathogenicity [29, 35]. According to the ACMG guidelines for reclassification of variants, ATPase activity data from functional analysis provide strong evidence for the pathogenic strong-3 (PS3) and benign strong-3 (BS3) categories [35]. Functional data derived from assays that more closely resemble the actual biological environment of the variants are required for reclassification. Bacterial expression of recombinant ABCA4 variants, followed by functional assays, has also been reported in the literature; however, this approach suffers from protein insolubility and does not recapitulate the native protein environment [36, 37]. The baculovirus expression system employed in this study generates membrane-localized recombinant variant proteins in VLPs. This approach reflects the biological milieu, lipid environment and the topology of the ABCA4 protein in the photoreceptor disc membrane and provides additional information about the membrane-localization potential of individual VUS. Additionally, this approach also permits essential post-translational modifications of ABCA4, including N-linked glycosylation.

Despite the variations in expression and localization, p.R140P, p.R140W, p.R140Q, p.H193Y, p.D197N, and p.N269H variants showed significant NRPE-stimulated ATPase activities compared to their respective basal ATPase activities (Fig 7). But their NRPE-stimulated ATPase activities were reduced by ≥20% that of the NRPE-stimulated ATPase activity of the WT. Therefore, these VUS might lead to a later onset of IRDs due to reduced efficiency in retinoid transport [38]. At the multiallelic site p.R140, the VUS p.R140P is associated with retinal dystrophy, according to ClinVar, and a complex allele containing p.R140Q has been found in a 20-year-old patient with STGD1 symptoms, including flecks in the RPE suggestive of a metabolic deposit and distorted RPE [34]. For the reclassification of the above VUS at p.R140, p.H193Y, p.D197N, and p.N269H, reduced ATPase activities from the functional assay provide evidence to support PS3_supporting criteria. Additionally, the low allele frequency (<0.0001) of these VUS (Table S1) in population databases support the criteria PM2_supporting [35, 39].

**Fig 7.**
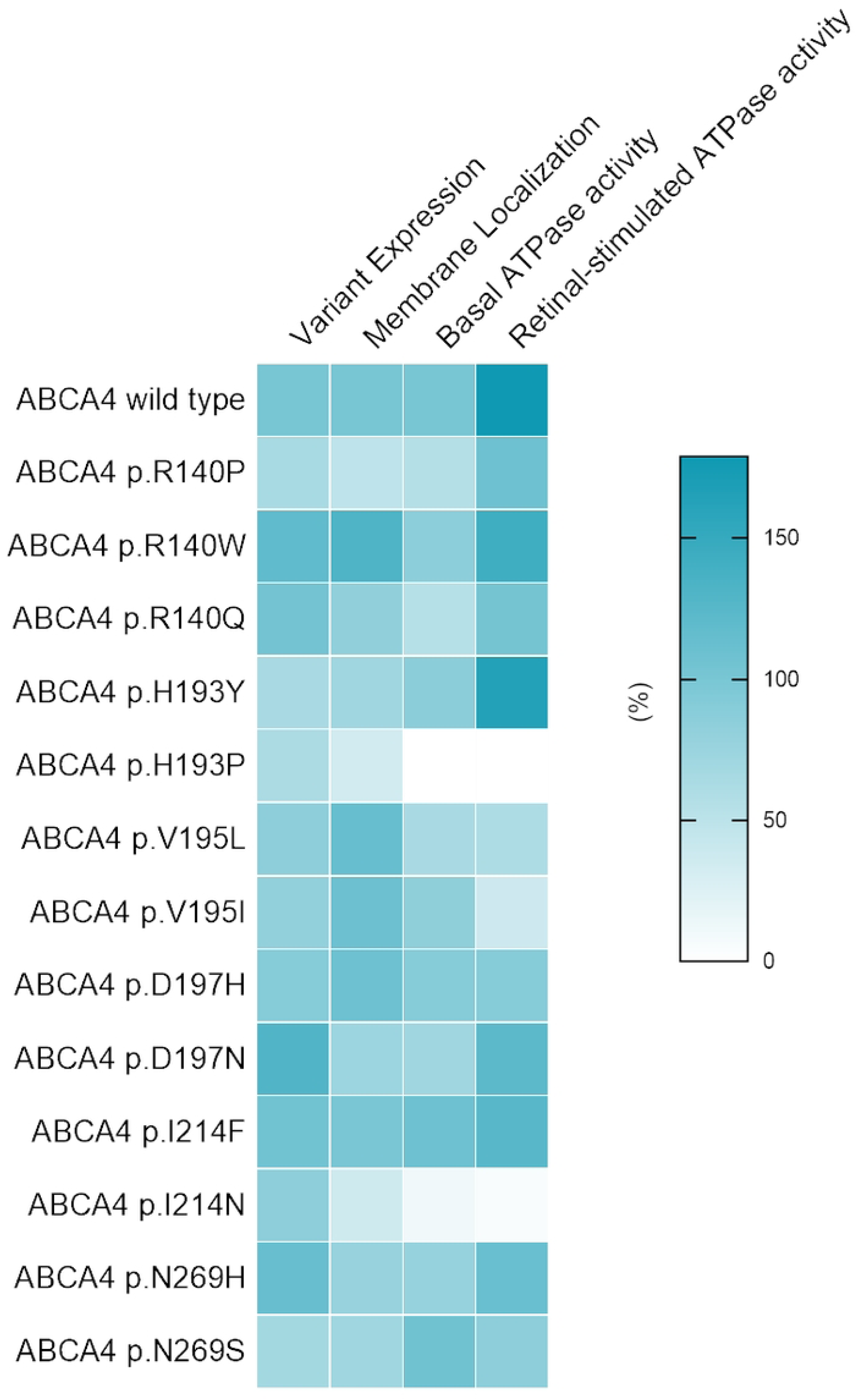
A summary of the functional analysis of the ECD1 uncharacterized region missense VUS. The variant expression in Hi5 cells and its localization to the VLP membrane were quantified by western blot analysis and normalized to WT levels (100%). The basal and retinal-stimulated ATPase activities are from the ATPase activity assay conducted using 1 ug of the VLP in the presence and the absence of retinal substrate NRPE. The ATPase activities were normalized based on the basal ATPase activity (100%) of the WT.

Missense VUS p.V195L, p.V195I, p.D197H, p.I214F, and p.N269S did not exhibit NRPE-stimulated ATPase activities despite proper membrane localization, similar to wild type (Fig 7). Therefore, these variants are functionally impaired and exhibit a strong pathogenic feature. Despite the conserved polarity of the p.V195L and p.V195I variants, they are associated with retinal dystrophy, and p.I214F is linked to an unspecified inherited genetic disease, further supporting their disease association in the clinical context. The lack of retinal stimulation of the VUS at p.V195, p.D197H, p.I214F and p.N269S explicitly demonstrates a functional deficit under PS3_ supporting criteria. Furthermore, the absence or low allele frequencies of VUS p.V195L, p.V195I, p.I214F, and p.N269S (Table S1) provide evidence for pathogenicity under the criteria PM2_supporting. However, the VUS p.D197H shows an allele frequency >0.0001 (Table S1) and does not meet PM2_supporting criteria.

Structurally unstable and misfolding variants p.H193P and p.I214N exhibited loss of ATPase activity, consistent with their predicted pathogenicity in the *in silico* platform and were not reported to be associated with disease (Fig 7). Their reclassifications are supported by PS3_supporting, PM2_supporting, and PP3_supporting criteria [35].

Functional studies conducted on VUS-containing animal samples, retinal organoids, or induced pluripotent stem cells (iPSCs) would further strengthen the evidence disclosed in this study. Although NRPE translocation by ABCA4 is mechanistically coupled to ATP hydrolysis, functional analysis based on the ATPase activity of ABCA4 variants is an indirect assessment of substrate transport. Therefore, direct evaluation of NRPE transfer across these variants would yield more robust evidence to support functional characterization and reclassification.

Apart from the reclassification of these missense VUS in the uncharacterized region, this study provides mechanistic insights into the potential functional roles of the ECDs. There are several proposed functional roles for ECD domains of ABCA4. They are speculated to have a regulatory role in retinoid transport across the disc membrane [10]. Although evidence is scarce, ECDs are also thought to interact with small molecules or proteins in the disc lumen via their large, flexible loops [10]. Certain prokaryotic ABC transporters interact with other luminal proteins, which act as substrate receptors to access their substrates [40]. It remains to be proven whether ABCA4 ECDs, especially the lid and tunnel, play such a role in accessing NRPE.

Although ABCA4 cryo-EM structures provide ample information for understanding ATP-driven NRPE translocation through ABCA4, the involvement of ECD1 remains to be elucidated. In the apo state, the two TMDs of ABCA4 are in an outward-facing conformation. The major retinoid substrate, NRPE, binds to a specific preformed binding site at the lumen leaflet of the disc membrane surrounded by TMD1, TMD2, and capped by an S-loop from ECD1 [10]. The phosphate group of NRPE is stabilized by electrostatic interactions with two arginine residues: p.R653 in TMD1 and p.R587 in ECD1 [10]. The fatty acyl chain was stabilized by multiple nonpolar residues from TMDs and ECD1 [10]. Given that, affected retinal stimulation of non-polar variants (p.V195L, p.V195I, p.I214F) might arise from disrupted NRPE-binding interactions at the lid of ABCA4. ECD1 is the largest domain of ABCA4, consisting of nearly 600 amino acids and ClinVar identifies approximately 17 pathogenic and likely pathogenic variant sites in the uncharacterized region of ECD1 (from residue 138 to 271), out of which 11 are non-polar residues, mostly mutating to polar residues [10, 13]. Furthermore, ECD1 has the largest proportion of missense VUS (n=236, 27%) among all ABCA4 domains, and 52 missense VUS map to the uncharacterized region of ABCA4 (Table S2). Among them, 16 were predicted to be likely pathogenic by the *in silico* platform in this study, and 10 of these were non-polar residues mutating to polar residues. This underscores the importance of nonpolar residues in this uncharacterized lid domain. Lipid-like molecules were detected in the ECD1 hydrophobic tunnel region of the cryo-EM structures, and the lid could engage in similar ligand-binding interactions with its non-polar residues, modulating ABCA4 structure and function [10, 14, 15, 41].

The literature suggests a lateral-access mechanism for the NRPE substrate to access ABCA4. According to this proposed model, NRPE accesses the binding site through the membrane due to its solubility in the phospholipids of the membrane, bypassing the ECDs. On the contrary, ABCA1, which is the structurally closest family member of ABCA4, also provides insights for the substrate transport mechanism of this ABCA family. ABCA1 is highly homologous to ABCA4, sharing around 50% overall sequence identity and 24% sequence identity in the ECDs [10]. ABCA1 is known for its role in cholesterol efflux from the cytoplasm to Apo A-I in the extracellular environment, forming nHDL in the liver, intestine, and macrophages [41]. In ABCA1 cryo-EM structures, cholesterol-like densities were observed in the ECD1 lid region, and cholesterol is thought to escape to the extracellular environment through the lid-tunnel region of the ECD1 [41]. Given the conserved sequence and organization between ABCA1 and ABCA4 ECDs, it is reasonable to assume that the ABCA4 lid-tunnel has a similar function. Furthermore, the lid region of ABCA4 is more elongated than that of ABCA1, which might be an adaptation to accommodate relatively larger lipids/lipid-like molecules. The disease association of non-polar missense VUS (p.V195L, p.V195I, and p.I214F) with selectively impaired NRPE-stimulated activity suggests that the affected residues may participate, directly or indirectly, in interactions with the NRPE substrate, indicating a potential NRPE-access pathway. However, identification of NRPE or other small-molecule binding in this region remains challenging due to structural ambiguity within the uncharacterized lid region in available cryo-EM structures.

The pathogenicity of different VUS at a particular multiallelic site is primarily determined by their contributions to overall structural stability and substrate engagement. Additional robust experimentation on the structure and intermolecular interactions of the ECD1 domain is required to elucidate its role in the retinoid transfer and subsequent pathogenic mechanisms. It is apparent that in vitro expression and localization profiles, as well as functional analyses, outweigh *in silico* predictions, especially for characterizing variants in structurally and functionally uncharacterized regions; however, an integrated analysis would be more rigorous for VUS in more well-defined regions of ABCA4. The integrated analysis of the ABCA4 missense VUS at multiallelic sites in the uncharacterized region provides a comprehensive evaluation and information to reclassify these VUS into defined categories, such as pathogenic, likely pathogenic, benign, and likely benign, in accordance with ACMG guidelines. The reclassification of a VUS is a significant milestone for the community related to STGD1. It allows a number of STGD1 patients with such VUS to begin treatment at an early stage of the disease through clinical trials. Furthermore, reclassifying ABCA4 VUS meaningfully enhances genetic counseling for STGD1 by providing clearer insights into disease risk and progression. In this way, VUS reclassification strengthens genetic counseling as a more practical and forward-looking component of patient care.

## Acknowledgements

The authors gratefully acknowledge the Delaware Biotechnology Institute (DBI) and the Department of Medical and Molecular Sciences at the University of Delaware for providing access to shared instrumentation and research facilities that supported this study. We are grateful to the previous Biswas-Fiss lab researchers who laid the foundation for this work, including Dr. Senem Cevik.

## Supporting information

**S1 Fig. The structural conformation of the ECDs of ABCA4. A)** The representation of the lid-tunnel-base conformation of the ABCA4 ECDs derived from the cryo-EM structure [13] (PDB:7LKP) B) The AlphaFold 3 modeled structural conformation of the ECD1 of ABCA4. The ABCA4 ECD1 amino acid sequence from residues 45-648 was used to predict the secondary structure of the uncharacterized region of ECD1. The per-residue confidence level (pLDDT) of the modeling is represented in different colors in the structure. Dark blue: Very high (pLDDT > 90), cyan: High (90 > pLDDT > 70), yellow: Low (70 > pLDDT > 50) and orange: Very low (pLDDT < 50).

**S1 Table. Summary of the structural and functional effects of the 13 missense VUS and their corresponding ACMG criteria [39].**

**S2 Table. In Silico pathogenicity prediction scores and predicted pathogenicity of the 52 missense VUS in the uncharacterized region of the ECD1 domain of ABCA4**

## References

1. Ben-Yosef T. Inherited Retinal Diseases. Int J Mol Sci. 2022;23(21).

2. Hanany M, Rivolta C, Sharon D. Worldwide carrier frequency and genetic prevalence of autosomal recessive inherited retinal diseases. Proc Natl Acad Sci U S A. 2020;117(5):2710–6.

3. Praidou A, Hagan R, Newman W, Chandna A. Early diagnosis of Stargardt disease with multifocal electroretinogram in children. Int Ophthalmol. 2014;34(3):613–21.

4. Tanna P, Strauss RW, Fujinami K, Michaelides M. Stargardt disease: clinical features, molecular genetics, animal models and therapeutic options. Br J Ophthalmol. 2017;101(1):25–30.

5. Corradetti G, Verma A, Tojjar J, Almidani L, Oncel D, Emamverdi M, et al. Retinal Imaging Findings in Inherited Retinal Diseases. J Clin Med. 2024;13(7).

6. Crouch RK, Koutalos Y, Kono M, Schey K, Ablonczy Z. A2E and Lipofuscin. Prog Mol Biol Transl Sci. 2015;134:449–63.

7. Al-Khuzaei S, Broadgate S, Foster CR, Shah M, Yu J, Downes SM, et al. An Overview of the Genetics of ABCA4 Retinopathies, an Evolving Story. Genes (Basel). 2021;12(8).

8. Sun H, Nathans J. Mechanistic studies of ABCR, the ABC transporter in photoreceptor outer segments responsible for autosomal recessive Stargardt disease. J Bioenerg Biomembr. 2001;33(6):523–30.

9. Sun H, Smallwood PM, Nathans J. Biochemical defects in ABCR protein variants associated with human retinopathies. Nat Genet. 2000;26(2):242–6.

10. Scortecci JF, Molday LL, Curtis SB, Garces FA, Panwar P, Van Petegem F, et al. Cryo-EM structures of the ABCA4 importer reveal mechanisms underlying substrate binding and Stargardt disease. Nat Commun. 2021;12(1):5902.

11. Molday RS. ATP-binding cassette transporter ABCA4: molecular properties and role in vision and macular degeneration. J Bioenerg Biomembr. 2007;39(5-6):507–17.

12. Molday RS, Garces FA, Scortecci JF, Molday LL. Structure and function of ABCA4 and its role in the visual cycle and Stargardt macular degeneration. Prog Retin Eye Res. 2022;89:101036.

13. Liu F, Lee J, Chen J. Molecular structures of the eukaryotic retinal importer ABCA4. Elife. 2021;10.

14. Beharry S, Zhong M, Molday RS. N-retinylidene-phosphatidylethanolamine is the preferred retinoid substrate for the photoreceptor-specific ABC transporter ABCA4 (ABCR). J Biol Chem. 2004;279(52):53972–9.

15. Quazi F, Lenevich S, Molday RS. ABCA4 is an N-retinylidene-phosphatidylethanolamine and phosphatidylethanolamine importer. Nat Commun. 2012;3:925.

16. Cevik S, Jones JS, Biswas SB, Biswas-Fiss EE. Deciphering the impact of ABCA4 genetic variants of unknown significance in inherited retinal disease through computational and functional approaches. Adv Protein Chem Struct Biol. 2025;147:423–60.

17. Cevik S, Biswas SB, Biswas-Fiss EE. Assessment of ABCA4 Genetic Variants: Current Landscape and Future Prospects. Adv Exp Med Biol. 2025;1468:63–7.

18. Marie M, Bigot K, Angebault C, Barrau C, Gondouin P, Pagan D, et al. Light action spectrum on oxidative stress and mitochondrial damage in A2E-loaded retinal pigment epithelium cells. Cell Death Dis. 2018;9(3):287.

19. Cevik S, Biswas SB, Biswas-Fiss EE. Structural and Pathogenic Impacts of ABCA4 Variants in Retinal Degenerations-An In-Silico Study. Int J Mol Sci. 2023;24(8).

20. Garces F, Jiang K, Molday LL, Stohr H, Weber BH, Lyons CJ, et al. Correlating the Expression and Functional Activity of ABCA4 Disease Variants With the Phenotype of Patients With Stargardt Disease. Invest Ophthalmol Vis Sci. 2018;59(6):2305–15.

21. Lewis RA, Shroyer NF, Singh N, Allikmets R, Hutchinson A, Li Y, et al. Genotype/Phenotype analysis of a photoreceptor-specific ATP-binding cassette transporter gene, ABCR, in Stargardt disease. Am J Hum Genet. 1999;64(2):422–34.

22. Cevik S, Biswas SB, Ghosh A, Biswas-Fiss EE. Virus-like particles as robust tools for functional assessment: Deciphering the pathogenicity of ABCA4 genetic variants of uncertain significance. J Biol Chem. 2024;300(10):107739.

23. Aslaksen S, Aukrust I, Molday L, Holtan JP, Jansson RW, Berland S, et al. Functional Characterization of ABCA4 Missense Variants Aids Variant Interpretation and Phenotype Prediction in Patients With ABCA4-Retinal Dystrophies. Invest Ophthalmol Vis Sci. 2024;65(10):2.

24. Xie T, Zhang Z, Fang Q, Du B, Gong X. Structural basis of substrate recognition and translocation by human ABCA4. Nat Commun. 2021;12(1):3853.

25. Tsybovsky Y, Molday RS, Palczewski K. The ATP-binding cassette transporter ABCA4: structural and functional properties and role in retinal disease. Adv Exp Med Biol. 2010;703:105–25.

26. Abramson J, Adler J, Dunger J, Evans R, Green T, Pritzel A, et al. Accurate structure prediction of biomolecular interactions with AlphaFold 3. Nature. 2024;630(8016):493–500.

27. Krieger E, Vriend G. YASARA View - molecular graphics for all devices - from smartphones to workstations. Bioinformatics. 2014;30(20):2981–2.

28. Schymkowitz J, Borg J, Stricher F, Nys R, Rousseau F, Serrano L. The FoldX web server: an online force field. Nucleic Acids Res. 2005;33(Web Server issue):W382–8.

29. Ioannidis NM, Rothstein JH, Pejaver V, Middha S, McDonnell SK, Baheti S, et al. REVEL: An Ensemble Method for Predicting the Pathogenicity of Rare Missense Variants. Am J Hum Genet. 2016;99(4):877–85.

30. Porretta AP, Fressart V, Surget E, Morgat C, Bloch A, Messali A, et al. Making Sense of Missense: Benchmarking MutScore for Variant Interpretation in Inherited Cardiac Diseases. Mol Diagn Ther. 2025;29(4):539–52.

31. Li C, Zhi D, Wang K, Liu X. MetaRNN: differentiating rare pathogenic and rare benign missense SNVs and InDels using deep learning. Genome Med. 2022;14(1):115.

32. Bradford MM. A rapid and sensitive method for the quantitation of microgram quantities of protein utilizing the principle of protein-dye binding. Anal Biochem. 1976;72:248–54.

33. Biswas-Fiss EE. Functional analysis of genetic mutations in nucleotide binding domain 2 of the human retina specific ABC transporter. Biochemistry. 2003;42(36):10683–96.

34. Xiao X, Ye L, Chen C, Zheng H, Yuan J. Clinical Observation and Genotype-Phenotype Analysis of ABCA4- Related Hereditary Retinal Degeneration before Gene Therapy. Curr Gene Ther. 2022;22(4):342–51.

35. Richards S, Aziz N, Bale S, Bick D, Das S, Gastier-Foster J, et al. Standards and guidelines for the interpretation of sequence variants: a joint consensus recommendation of the American College of Medical Genetics and Genomics and the Association for Molecular Pathology. Genet Med. 2015;17(5):405–24.

36. Biswas-Fiss EE, Kurpad DS, Joshi K, Biswas SB. Interaction of extracellular domain 2 of the human retina-specific ATP-binding cassette transporter (ABCA4) with all-trans-retinal. J Biol Chem. 2010;285(25):19372–83.

37. Jones JS, Biswas SB, Biswas-Fiss EE. Exploring the Role of ABCA4’s ECD2 Domain in Inherited Retinal Degeneration: Computational and Functional Perspectives. Adv Exp Med Biol. 2025;1468:69–74.

38. Mihalek I, De Bruyn H, Glavan T, Lancos AM, Ciolfi CM, Malendowicz K, et al. Quantifying the Progression of Stargardt Disease in Double-Null ABCA4 Carriers Using Fundus Autofluorescence Imaging. Transl Vis Sci Technol. 2025;14(3):16.

39. Cornelis SS, Bauwens M, Haer-Wigman L, De Bruyne M, Pantrangi M, De Baere E, et al. Compendium of Clinical Variant Classification for 2,246 Unique ABCA4 Variants to Clarify Variant Pathogenicity in Stargardt Disease Using a Modified ACMG/AMP Framework. Hum Mutat. 2023;2023:6815504.

40. Davidson AL, Dassa E, Orelle C, Chen J. Structure, function, and evolution of bacterial ATP-binding cassette systems. Microbiol Mol Biol Rev. 2008;72(2):317–64.

41. Chen L, Zhao ZW, Zeng PH, Zhou YJ, Yin WJ. Molecular mechanisms for ABCA1-mediated cholesterol efflux. Cell Cycle. 2022;21(11):1121–39.

